# Intracellular *Burkholderia* Symbionts Induce Extracellular Secondary Infections; Driving Diverse Host Outcomes that Vary by Genotype and Environment

**DOI:** 10.1101/448399

**Authors:** Niloufar Khojandi, Tamara S. Haselkorn, Madison N. Eschbach, Rana A. Naser, Susanne DiSalvo

## Abstract

Symbiotic associations impact and are impacted by their surrounding ecosystem. The association between *Burkholderia* bacteria and the soil amoeba *Dictyostelium discoideum* is a tractable model to unravel the biology underlying symbiont-endowed phenotypes and their impacts. Several *Burkholderia* species stably associate with *D. discoideum* and typically reduce host fitness in food-rich environments while increasing fitness in food-scarce environments. *Burkholderia* symbionts are themselves inedible to their hosts but induce co-infections with secondary bacteria that can serve as a food source. Thus, *Burkholderia* hosts are “farmers” that carry food bacteria to new environments, providing a benefit when food is scarce. We examined the ability of specific *Burkholderia* genotypes to induce secondary co-infections and assessed host fitness under a range of co-infection conditions and environmental contexts. Although all *Burkholderia* symbionts intracellularly infected *Dictyostelium,* we found that co-infections are predominantly extracellular, suggesting that farming benefits are derived from extracellular infection of host structures. Furthermore, levels of secondary infection are linked to conditional host fitness; *B. agricolaris* infected hosts have the highest level of co-infection and have the highest fitness in food scarce environments. This study illuminates the phenomenon of co-infection induction across *Dictyostelium* associated *Burkholderia* species and exemplifies the contextual complexity of these associations.

## Introduction

Symbiotic interactions can alter the fitness and evolutionary trajectory of both partners (1–4). Clearly detrimental or mutualistic associations have been investigated for obvious reasons: to eliminate infectious disease, boost health, and restore ecosystems. However, many symbiotic associations evade simple characterization and related mechanisms can underlie opposing outcomes (5,6). Invasion and replication strategies employed by mutualists and pathogens often resemble each other, while genotypes and external factors modify subsequent outcomes (7). Genotype pairing determines the outcome of plant-mycorrhizae interactions (8) and amplification of a genomic region in a normally beneficial *Wolbachia* symbiont leads to over-replication at the hosts expense (9). Light mediates pathogenicity of a fungal plant endosymbiont (10), temperature affects reproductive fitness of aphids hosting *Buchnera* (11), and parasitoid pressure determines whether *Hamiltonella defensa* is beneficial to host aphids (12). These examples demonstrate that even canonically beneficial or detrimental associations may produce alternative effects in alternative contexts (4,13–17).

Eukaryotic microbes, such as amoebae, are attractive models for exploring eukaryote-prokaryote interactions. Amoebae are ubiquitous and efficient phagocytic predators of bacterial prey, making them important shapers of the microbial community (18). This pressures prey microbes to evolve virulence strategies that enable evasion of phagocytosis or subsequent digestion (19). Amoebae are thereby potential training grounds and environmental reservoirs for bacterial pathogens. Amoebae phagocytosis also enables bacteria to gain easy access to an attractive intracellular niche, bypassing the requirement for evolving specialized cell-entry mechanisms. After invasion, bacteria can be retained in an environmentally resistant cyst or spore (20). A number of bacterial pathogens, such as *Legionellae pneumophila* and others (21,22), are harbored in different species of amoebae and there is a growing list of recently identified amoebae symbionts (23,24).

The social amoebae *Dictyostelium discoideum* has been appreciated as a model host for studying bacterial pathogens for some time (25–27). Recently, work with wild isolates has emphasized its power for exploring naturally occurring microbial symbioses (28,29). As a social amoeba *Dictyostelium* exhibits a unique life cycle, transitioning between single-and multi-cellular forms. Under favorable conditions, it lives as a unicellular amoeba, consuming bacteria and dividing by binary fission. When bacterial food is depleted, amoebae secrete cAMP, which triggers the transition to multi-cellularity. During this phase, amoebae aggregate to form a multicellular slug that seeks out a location for fruiting body formation (such as the soil surface). Fruiting bodies are comprised of a spherical sorus containing hardy spore cells resting atop a long stalk of dead cells. This positioning of spore cells likely aids in their dispersal (30). Once dispersed, spores germinate, and the cycle continues.

*D. discoideum* grown with a variety of bacterial food traditionally form germ-free sori, clearing residual bacteria from the multicellular state during development. Microbial clearance is aided by immune-like sentinel cells which engulf debris and slough off the migrating slug (31,32). However, approximately one third of wild *D. discoideum* isolates are naturally and stably colonized by *Burkholderia* bacteria (33,34). *Burkholderia* can be easily eliminated from host populations with antibiotic treatment and new associations can be readily initiated through co-culture. These *Burkholderia* symbionts establish intracellular infections which persist through host development, resulting in sori containing both extracellular and intracellular bacteria (34). *Burkholderia* symbionts thereby remain associated with host populations during spore dispersal and can be acquired through vertical and horizontal transmission routes. This mixed mode of transmission has interesting implications for the fitness consequences and evolutionary trajectory of the symbiosis.

*Burkholderia* symbionts of *D. discoideum* are members of the plant beneficial environmental group within the *Burkholderia* genus (35). Symbiont strains are genetically diverse, belonging to three species arising from two independent lineages: *B. agricolaris, B. hayleyella,* and *B. bonniea* (36). *Burkholderia* differentially impacts host fitness according to host-symbiont genotype combinations and environmental context (33,34,37). Symbionts generally reduce host fitness in food rich conditions but enhance fitness in food scarce environments (33,34,37). Fitness benefits are attributed to retention of bacteria within host spores, allowing them to reseed new environments with bacterial food. This trait is called farming and *Burkholderia* infected hosts are thus referred to as “farmers”. *Burkholderia* symbionts themselves are poor food sources for their hosts (33,34). However, *Burkholderia* infection appears to increase host susceptibility to secondary bacterial infection, promoting the formation of a mini-microbiome. It is these secondary bacteria that can serve as an amoebae food source and thereby provide the farming benefit.

Given the importance of secondary infection in farming, we sought to explore the underlying dynamics of this interaction. While a commonly used lab food strain, *K. pneumoniae,* can be identified as an occasional co-infecting partner, co-infection dynamics might vary depending on particular bacterial pairings (34). Thus, food bacteria identity is an important environmental context that may affect outcomes. Furthermore, the three different *Burkholderia* symbiont species have divergent evolutionary histories of association with *D. discoideum.* While they have converged on the farming phenotype, the effects and underlying mechanisms of infection may differ across *Burkholderia* species (Haselkorn et al. submitted).

Here, we reveal the density and location of secondary co-infections induced by each *Burkholderia* species with a collection of secondary bacteria. Next, we clarified the downstream benefits of *Burkholderia* infection in varied food availability contexts and link these with host outcomes according to co-infection induction. Specifically, we analyzed co-infection patterns and host outcomes with a variety of secondary bacteria including: laboratory food *Klebsiella pneumoniae, Rhizobium* and *Serratia* isolates that naturally co-occur with *D. discoideum,* and *Agrobacterium tumefaciens* and *Pseudomonas aeruginosa* as pathogens that *D. discoideum* may encounter in nature. We found that all *Burkholderia* symbionts induce some degree of secondary infection in host sori but the density and location of secondary infections is dependent on *Burkholderia* genotype and secondary bacterial identity. Contrary to previous inference, secondary infections are predominantly extracellular with intracellular co-infections only readily visualized in *B. agricolaris* infected spores. Overall, *B. agricolaris* induces the highest density of combined co-infection resulting in a higher fitness benefit hosts in food scarce environments. *B. bonniea* and *B. hayleyella* induce lower levels of secondary co-infection but only *B. bonniea* provides significant host benefits under specific dispersal conditions. This work illuminates the interplay between symbiont genotypes and environmental context in mediating the expression and consequences of novel symbiont endowed phenotypes.

## Materials and Methods

### Bacterial Strains and Culturing

*Burkholderia* strains were isolated in the Queller Strassmann lab from *D. discoideum* hosts. *Klebsiella pneumoniae, Rhizobium* and *Serratia* originate from the Queller Strassmann lab with *Rhizobium* and *Serratia* isolated from wild *Dictyostelium* fruiting bodies grown directly from Virginia Mountain Lake Biological Station soil and 16s rRNA gene sequenced with universal primers for putative identification (genbank accession numbers MH997560 and MH997561). *Pseudomonas aeruginosa* PAO1-GFP (derived from ATCC15692) was provided by R. Fred Inglis (38). *Agrobacterium tumefaciens* (A136) was provided by Daniel Gage (reclassified as *Rhizobium radiobacter* 51350 in ATCC). We used GFP labeled secondary bacteria for all experiments, with the exception of *K. pneumoniae,* which is unlabeled when mixed with other bacteria. We grew all bacteria on SM/5 medium (Formedium: 2g Peptone, 0.2g yeast extract, 2g glucose, 1.9g KH_2_PO_4_, 1.3g K_2_HPO_4_.3H_2_0, 0.49g MgO_4_.anhydrous, 17g agar/liter) at room temperature. To prepare bacteria for culturing *Dictyostelium*, we suspended bacterial colonies from SM/5 medium into KK2 (2.2g KH2PO4 monobasic and 0.7g K2HPO4 dibasic/liter) and set to an OD_600nm_ of 1.5. For *K.pneumoniae*/secondary bacterial mixtures, we combined bacterial suspensions equally by volume. For *Burkholderia* infections, we added 5% by volume *Burkholderia-RFP* to indicated bacterial mixtures.

### Construction of Fluorescent Bacterial Strains

We generated RFP labeled *Burkholderia* by triparental mating with *E. coli* helper strain E1354 (pTNS3-asdEc) and *E. coli* donor strain E2072 (pmini-Tn7-gat-P1-rfp) and confirmed identity of RFP conjugants using a *Burkholderia* specific PCR as previously described (34,39,40). We GFP labeled *Rhizobium, Serratia, A. tumefaciens,* and *K. pneumoniae* through triparental mating with *E. coli* donor WM3064 (pmini-Tn7-KS-GFP) and *E. coli* helper E1354 (pUXBF13) as previously described (41) and confirmed identity of GFP positive conjugants through 16s rRNA gene sequencing. *P. aeruginosa*-GFP was described previously (38).

### Dictyostelium Culture Conditions

We used *D. discoideum* clone QS864 (naturally symbiont free) for all experiments. Cultures were initiated by plating spores on SM/5 medium with *K. pneumoniae* and incubating under lights at room temperature until fruiting bodies developed (4-7 days). For experiments, 10^5^ spores were harvested from developed sori and plated with 200μL of the appropriate bacterial mixtures. For all experiments (unless otherwise indicated) we analyzed sori 5 days after plating.

For co-infection assays we plated uninfected spores on bacterial mixtures with *Burkholderia,* uninfected controls were plated without *Burkholderia.* To compare spore productivity under food variable conditions, we harvested sori from indicated co-infection conditions and plated 10^5^ spores onto SM/5 with K. *pneumoniae* at an OD600 of 1.5 for food-rich conditions or with heat-killed (30 min at 80°^C^) K. *pneumoniae* at an OD_600_ of 6 for food scarce conditions.

### Spore Production Assays

To harvest total spores, we flooded each plate with 5-10 mL KK2 + 0.1% Nonidet P-40 alternative and collected the entire surface contents into 15-mL Falcon tubes. We then diluted samples in KK2 and counted spores on a hemocytometer. At least five replicates were analyzed for each treatment.

### Confocal Microscopy

We imaged spores by staining with 1% calcofluor in KK2, placing on glass bottom culture dishes (Electron Microscopy Sciences) and overlaying with 2% agarose. We imaged samples on an Olympus Fluoview FV1000 confocal microscope using Plan Apo Oil 1.4NA 60X objective. Z-sections were taken every 0.5 microns at 1024 resolution. Calcofluor was visualized with DAPI, GFP with FITC, and RFP with Cy3 then pseudocolored grey, green, and red respectively. We imaged at least three individual replicates and counted more than 30 spores for each.

### Colony forming unit quantification

To quantify secondary bacteria, we harvested sori grown from the indicated co-culture conditions from 6-or 14-day incubations. We suspended individual sori in KK2+0.05% Nonidet P-40 alternative, counted spores on a hemocytometer, plated serial dilutions on SM/5 medium and incubated plates at room temperature until colony formation (∼2 days), and counted GFP colonies using a safe-light imaging system. We performed 3 or more independent replicates for each treatment.

### Streak test

Our streak test assay was initiated from the indicated co-culture conditions by touching individual sori with sterile pipette tips and transferring them to SM/5 plates along a ∼1 inch streak. We incubated plates face up under lights at room temperature and examined them 5 days (or 2 weeks) after streaking. We determined the percentage of streaks with bacterial growth, percentage of bacterial positive streaks with fruiting bodies, and number of fruiting bodies in positive streaks. Streaks were photographed on a Cannon Eos7D with a macro-lens. Six sori were streaked for each replicate for at least 4 individual replicates per condition.

### Statistical analysis

We analyzed all data using R (version 3.3.1). For normally distributed data we determined significance using a standard one-way analysis of variance (ANOVA) and a *post hoc* Tukey HSD test. For non-normally distributed data we performed a Kruskall-wallis test and *post hoc* analysis with a Dunn test using the dunnTest function in the FSA package. We used *Burkholderia* status as fixed effects for all conditions.

## Results

### *Burkholderia* and Secondary Bacterial Combinations

To investigate the induction of secondary infection by *Burkholderia* symbionts, we cultured an uninfected natural isolate of *D. discoideum* with different *Burkholderia-RFP* and secondary bacteria-GFP combinations. We began with three *Burkholderia* strains: Ba.70, Bh.11, and Bb.859, each representing one *D. discoideum* symbiont species *B. agricolaris, B. hayleyella,* and *B. bonniea* (36). Secondary bacteria consisted of a *Klebsiella pneumoniae* strain, soil isolated *Rhizobium* and *Serratia,* and lab *Agrobacterium tumefaciens* and *Pseudomonas aeruginosa* strains. We chose these representatives because: 1) *K. pneumoniae* is used widely as a lab food source for *D. discoideum* and serves as a good starting point for experimental conditions. 2) The *Rhizobium* and *Serratia* strains where co-isolated with *D. discoideum* and are thereby ecologically relevant potential co-associates. 3) *A. tumefaciens,* in addition to its use in plant molecular biology, is an important soil dwelling plant pathogen. As such, amoebae may interact with *A. tumefaciens* in the environment and this could subsequently impact the surrounding ecosystem. 4) *P. aeruginosa* is an important opportunistic human pathogen whose association with other bacterial species in biofilms (such as pathogenic *Burkholderia cenocepacia*) influences infection outcomes (42,43). Adding *Pseudomonas* to the *Burkholderia-Dictyostelium* system provides a novel approach to explore microbial interactions and virulence.

### Host Outcomes Differ According to *Burkholderia* and Secondary Bacteria Conditions

First, we examined host fitness when amoebae were co-cultured with *Burkholderia* and secondary bacteria. We determined total spore productivity of host amoebae after one social cycle on each labeled secondary bacterium, either alone or in a 50% mixture with *K. pneumoniae.* Five percent by volume of *Burkholderia-RFP* was included to establish infections (Figure 1). *D. discoideum* was unable to develop on any conditions where *P. aeruginosa* was the only food source suggesting that this strain was toxic and/or inedible for amoebae. All other conditions supported fruiting body development, but spore productivity varied across conditions (Figure 1). In line with previous studies, *Burkholderia* species differentially impact spore productivity on *K. pneumoniae.* Typically, *B. hayleyella* was the most detrimental for host fitness with *B. agricolaris* and *B. bonniea* being neutral or moderately detrimental. However, these patterns and the degree by which symbiont altered host fitness varied across culture conditions (Figure 1 and Table 1). These results highlight the variability of fitness outcomes caused by distinct *Burkholderia* symbionts and suggest that surrounding bacterial communities also impact fitness outcomes.

**Figure 1.**
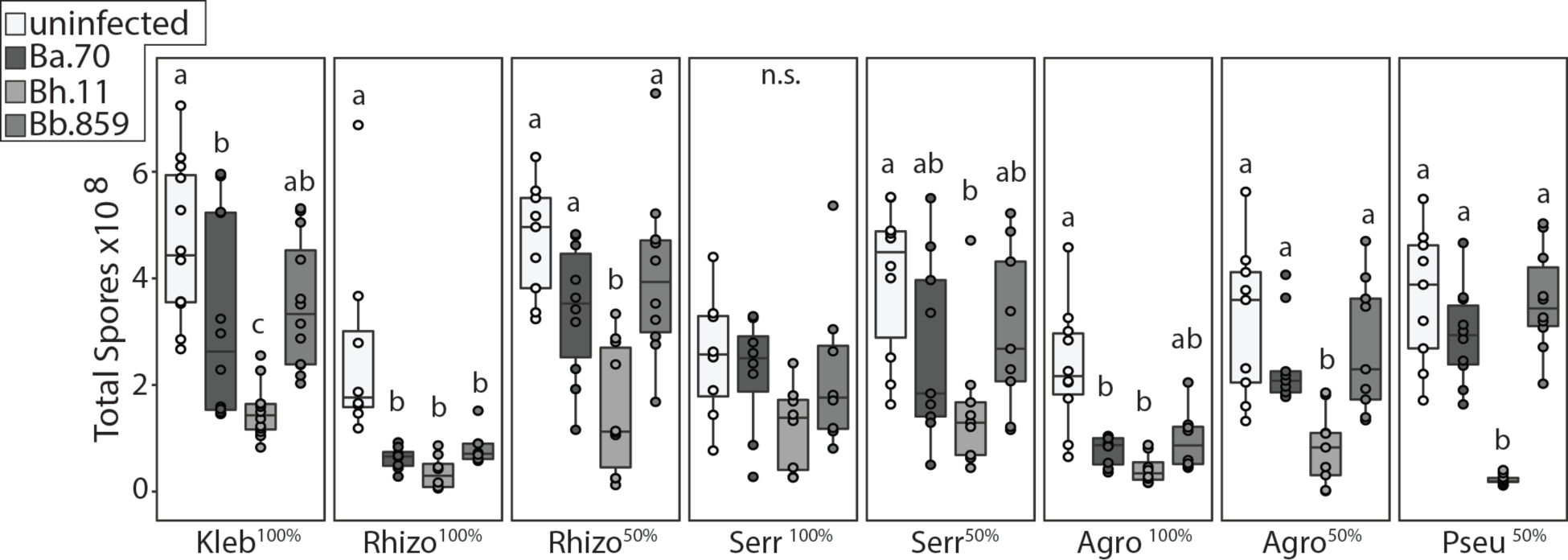
Culture Conditions Modify the Impact of *Burkholderia* Infections on Host Fitness. Total spore productivity for *D. discoideum* after one round of development on the indicated GFP labeled bacterial species (Kleb= *K. pneumoniae,* Rhizo= *Rhizobium,* Serr= *Serratia,* Agro=A, *tumefaciens,* and Pseu=P. *aeruginosa.* 50% cultures are mixed with 50% unlabeled *K. pneumoniae* by volume). Dark grey, light grey, and medium grey boxes indicate cultures wherein *Burkholderia* infections are initiated by inclusion of 5% *B. agricolaris.*70-RFP, *B. hayleyella.* 11-RFP, and *B. bonniea.*859-RFP, respectively. Points within boxes indicate individual replicates. Letters indicate post-hoc significance within panels.

**Table 1.** *Burkholderia* infections significantly alter spore productivity in most (but not all) bacterial culture conditions. Statistical Analysis of Spore Fitness from Figure 1. All DF values=3.

| Condition | $\chi^2$ | p value |
| --- | --- | --- |
| *K. pneumoniae*-GFP<sup>100%</sup> | 23.537 | >0.001 |
| *Rhizobium*-GFP<sup>100%</sup> | 20.81 | >0.001 |
| *Rhizobium*-GFP<sup>50%</sup> | 18.759 | >0.001 |
| *Serratia*-GFP<sup>100%</sup> | 6.411 | 0.09 |
| *Serratia*-GFP<sup>50%</sup> | 11.292 | 0.01 |
| *Agrobacterium*-GFP<sup>100%</sup> | 18.76 | >0.001 |
| *Agrobacterium*-GFP<sup>50%</sup> | 16.267 | >0.001 |
| *Pseudomonas*-GFP<sup>50%</sup> | 23.36 | >0.001 |

### Intracellular Co-infection is Rare and Depends on *Burkholderia* and Secondary Bacterial Combinations

To investigate the induction of secondary infections we imaged *D. discoideum* sori contents after development on *Burkholderia* and secondary bacteria. We used 50/50 *K. pneumoniae/secondary* bacteria-GFP conditions as they resulted in better amoebae development than secondary bacteria-only conditions. We also imaged sori grown from *K. pneumoniae-GFP.* Importantly, we do not detect any secondary bacteria in sori in the absence of *Burkholderia* (Figure 2). Thus, these bacteria are not capable of infecting *D. discoideum* on their own. In contrast, we can detect secondary-GFP cells in sori from amoebae co-exposed to *Burkholderia* (Figure 3). To determine their prevalence in host spore populations, we quantified the percent of spores intracellularly infected with *Burkholderia-RFP* and with secondary bacteria-GFP. First, the percent of spores infected with each *Burkholderia* species significantly differs (x^2^=44.02, df=2, p<0.001). On average across conditions, *B. hayleyella* infects the most (89.2%), *B. bonniea* infects an intermediate (68.6%), and *B. agricolaris* infects the fewest (33%) percent of spores. However, we only readily observe intracellular secondary co-infections in *B. agricolaris* host spores (average of 5.5% across conditions). Only rarely, if ever, do we observe intracellular GFP in *B. hayleyella* and *B. bonniea* hosts (0.01% and 0.05% respectively) (Figure 3). We did not observe intracellular GFP in the absence of intracellular RFP, suggesting that secondary bacteria are only retained in *Burkholderia* co-infected spores.

**Figure 2.**
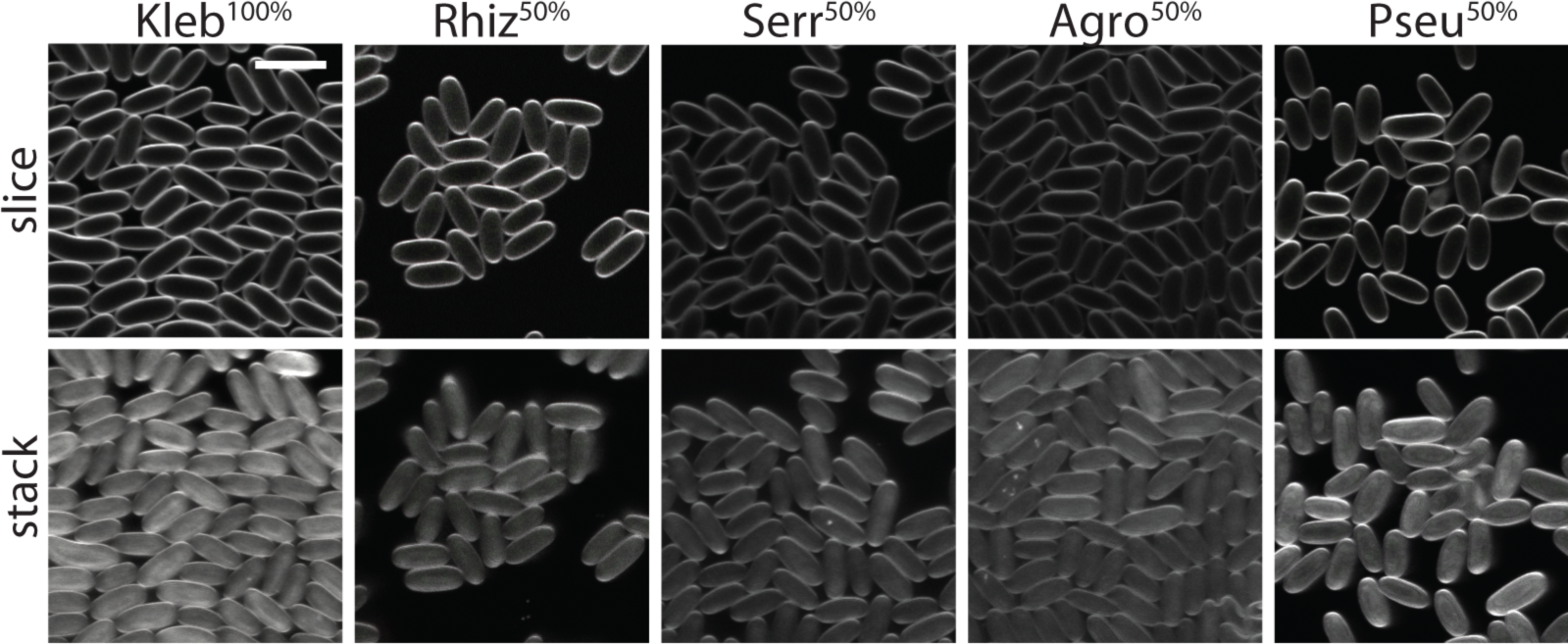
Secondary Bacteria do not Infect Symbiont Free Spores. Representative confocal micrographs of sori contents after growth on the indicated GFP labeled bacterial cultures. Scale bar=10um.

**Figure 3.**
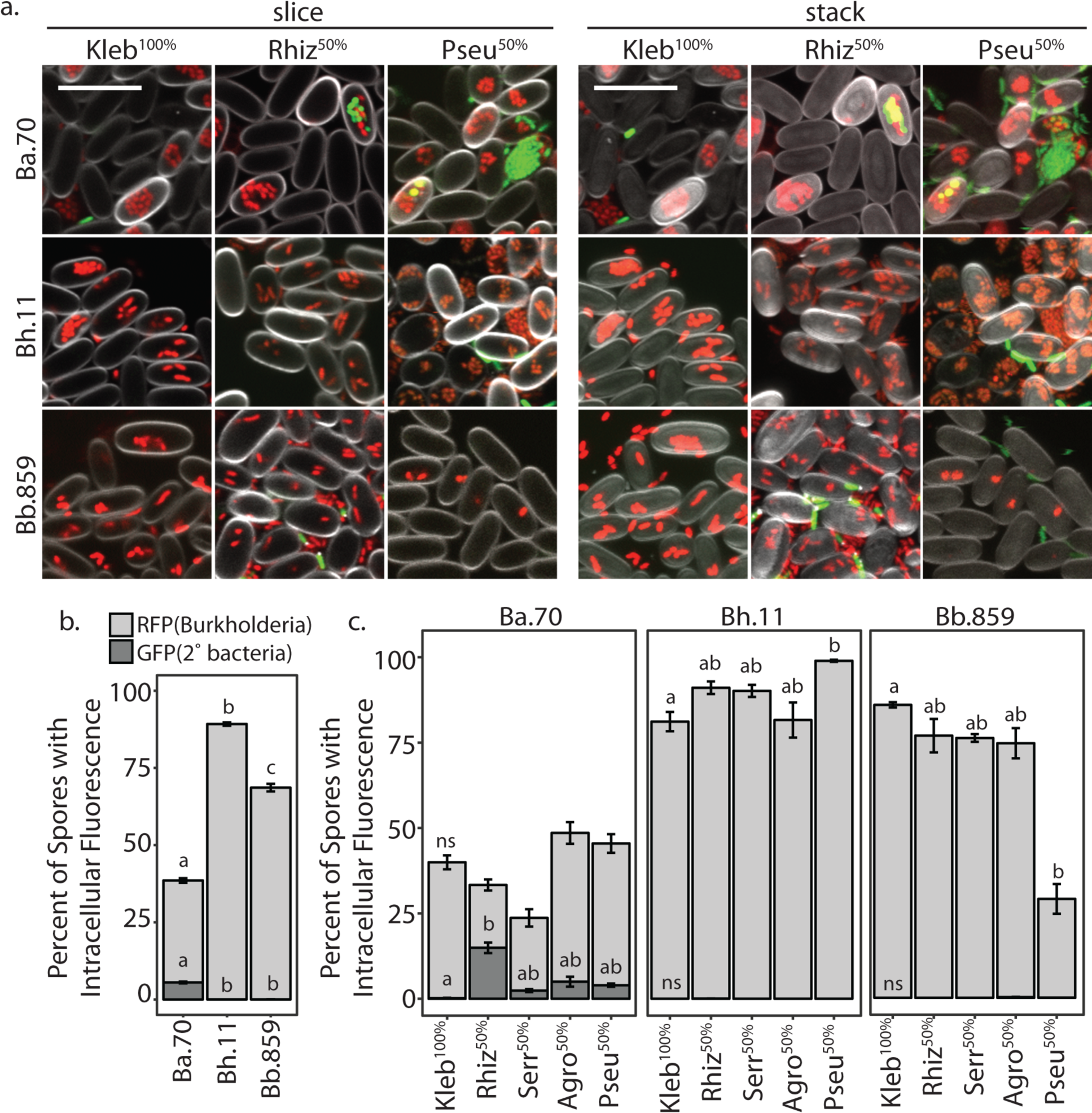
*Burkholderia* Differentially Induce Intracellular Co-Infections. **a)** Representative confocal micrographs of *Burkholderia-RFP* infected sori contents after development on the indicated GFP labeled secondary bacteria. Scale bar= 10um. b) Percent of spores infected with *Burkholderia-RFP* (light grey) and co-infected with secondary bacteria-GFP (dark grey) averaged across all replicates of all secondary bacterial conditions. c) Average percent of spores infected (as in b) for each secondary culture condition. Error bars ±SE.

The secondary bacterium also plays a role in the prevalence of intracellular infections for both *Burkholderia* and secondary bacteria. For instance, slightly fewer spores are infected with *B. bonniea* when cultured with *P.* aeruginosa-GFP (29%) than with all other bacteria (85.9-75.6%) (x^2^=10.843, df=4, p=0.028). On the flip side, significantly more spores are infected with *B. hayleyella* when cultured with *P.* aeruginosa-GFP (98.9%) than with all other bacteria (81.2-91%) (x^2^=10.217, df=4, p=0.036). For *B. agricolaris* hosts, the degree of secondary co-infections significantly varied by secondary bacteria (x^2^=15.019, df=4, p=0.004). *K. pneumoniae-GFP* is localized in only 0.2% of total spores while *Rhizobium-GFP* was localized in 14.9%. When secondary bacterial infections are considered as a percentage of spores co-infected with *B. agricolaris, Rhizobium-GFP* is co-localized in almost half of total infected spores. This suggests that should *B. agricolaris* infection levels increase due to a higher infectious dose or conditions that promote higher infection titers, secondary infections may correspondingly increase.

### *Burkholderia* Symbionts Induce Extracellular Secondary Infections

Although we found minimal intracellular co-infections in most conditions, the farming phenotype may instead by explained by extracellular secondary infections. To get an initial indication of extracellular co-infections, we determined the percent of confocal images in which any extracellular GFP could be visualized (Figure 4a). We found that all *Burkholderia* symbionts induced at least some level of extracellular co-infections, as we could visualize external GFP in confocal images for each condition. Similar to our observations for intracellular co-infections, extracellular secondary bacteria appeared most frequently in *B. agricolaris* host sori (Figure 4a).

**Figure 4.**
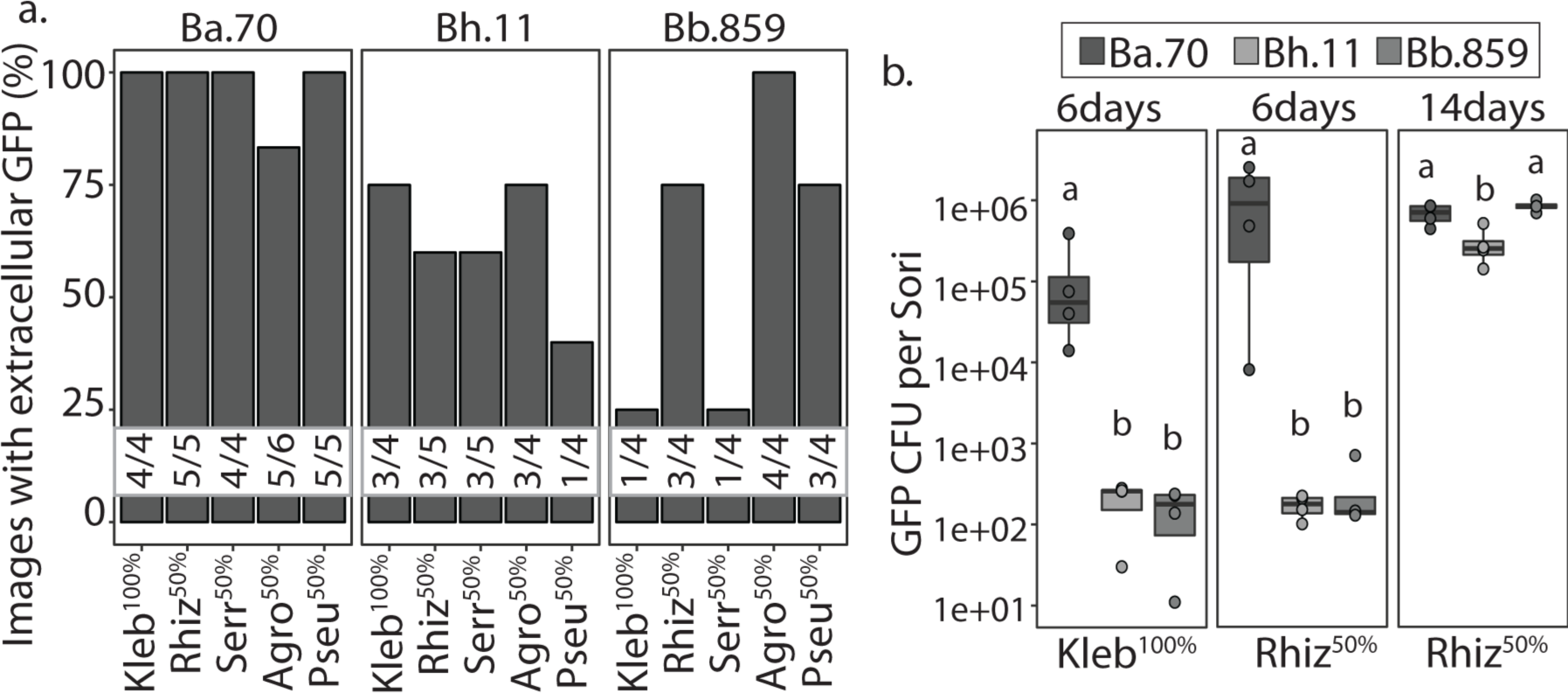
*Burkholderia* Induce Extracellular Co-infections. a) Percent of confocal micrographs wherein secondary bacterial-GFP cells were visualized extracellularly within sori contents. Numbers nested within bars indicate the number of images with visible GFP over the number of total images collected per condition. b) Number of GFP colony forming units (log 10) from *Burkholderia* infected sori contents when grown on *K. pneumoniae-GFP* and *Rhizobium-GFP.* Individual sori contents were harvested either 6 or 14 days after plating on the *Burkholderia/Secondary* bacterial culture conditions. These numbers may represent both extracellular and intracellularly derived secondary bacteria. Points represent the GFP-CFU count for individual replicates.

To quantify overall secondary co-infection, we counted GFP colony-forming units per sori for *K. pneumoniae*-GFP^100%^ and *Rhizobium*-GFP^50%^ conditions. After six days of co-culturing amoebae we counted the number of GFP positive colonies that developed from individual sori (Figure 4b). No bacterial colonies were recovered when we plated sori from fruiting bodies grown without *Burkholderia,* again indicating that these bacteria do not by themselves infect *D. discoideum.* We found that *B. agricolaris* induces the highest level of secondary infection, with *B. agricolaris* host sori colonized by an average of 1.29×10^5^ *K. pneumoniae-GFP* cfu’s and 1.17×10^6^ *Rhizobium-*GFP cfu’s (Figure 4b). We recover notable, albeit far fewer, secondary bacterial colonies from *B. hayleyella* and *B. bonniea* hosts. *B. hayleyella* sori host an average of 206 and 169, and *B. bonniea* an average of 153 and 276, *K. pneumoniae-GFP* and *Rhizobium-GFP* cfu’s, respectively.

To explore whether secondary bacteria could further amplify within fruiting bodies over time, we also quantified *Rhizobium-GFP* colony-forming units 14 days after plating. We found that cfu’s did not increase for *B. agricolaris* hosts, but dramatically increased for *B. hayleyella* and *B. bonniea* hosts, which produced 2.87×10^5^ and 8.37×10^5^ GFP cfu’s, respectively. This brought the number of cfu’s in all *Burkholderia* infected sori up to fairly similar levels, perhaps representing a peak carrying capacity. However, we noticed that the number of spores per sori for *B. hayleyella* and *B. bonniea* infected hosts appeared to decrease over time (not shown). Thus, replication of secondary bacteria within these sori could be damaging to spores, counter-acting potential benefits of hosting more food bacteria.

### *Burkholderia* Symbionts Benefit Hosts in Food Scarce Conditions in Relation to Coinfection Induction

Farmers have been shown to have an advantage when dispersed to food scarce environments (33,34). This is attributed to the induction of secondary bacterial food carriage, enabling host spores to reseed new environments with edible bacteria. However, this benefit has previously only been measured as an average fitness outcome across hosts infected with genotypically diverse *Burkholderia* symbionts (34). Whether or how specific *Burkholderia* genotypes correspond with this beneficial outcome remained unknown. Given our results demonstrating that *B. agricolaris* induces the highest level of secondary co-infections, we speculated that *B. agricolaris* hosts would have the highest reproductive fitness after dispersal to food scarce environments.

To compare host fitness under different food availability conditions, we first plated uninfected spores with *Burkholderia* and secondary bacteria (*K. pneumoniae*-GFP^100%^ and *Rhizobium*-GFP^50%^) under the same conditions employed previously. After five days of incubation we harvested developed sori and transferred 10^5^ spores with rich (live *K. pneumoniae-GFP*) or scarce (heat killed *K. pneumoniae)* food on nutrient medium. After five days of incubation in these conditions, we measured total spore productivity (Figure 5a). For food rich conditions, we again found significant differences according to amoebae infection status (F=7.41, df=3, p 0.0015 and x^2^=13.95, df=3, p=0.0029 for *K. pneumoniae-GFP*^100%^ and *Rhizobium*-GFP^50%^ conditions respectively). However, in this experiment only *B. hayleyella* hosts produced significantly fewer spores than the uninfected control (Figure 5a). For food scarce conditions, spore productivity was also significantly different according to *Burkholderia* infection status (x^2^=11.87 and 20.616, df=3, p=0.0078 and <0.001 for *K. pneumoniae*-GFP^100%^ and *Rhizobium*-GFP^50%^ conditions, respectively). Here, *B. agricolaris* hosts had the highest spore productivity for both secondary conditions (Figure 5a). *B. bonniea* also resulted in slightly higher, but not significantly different, spore productivity compared to the uninfected control (Figure 5a). Thus, *B. agricolaris* infections endow a benefit for their amoeba host when dispersed to food scarce environments.

**Figure 5.**
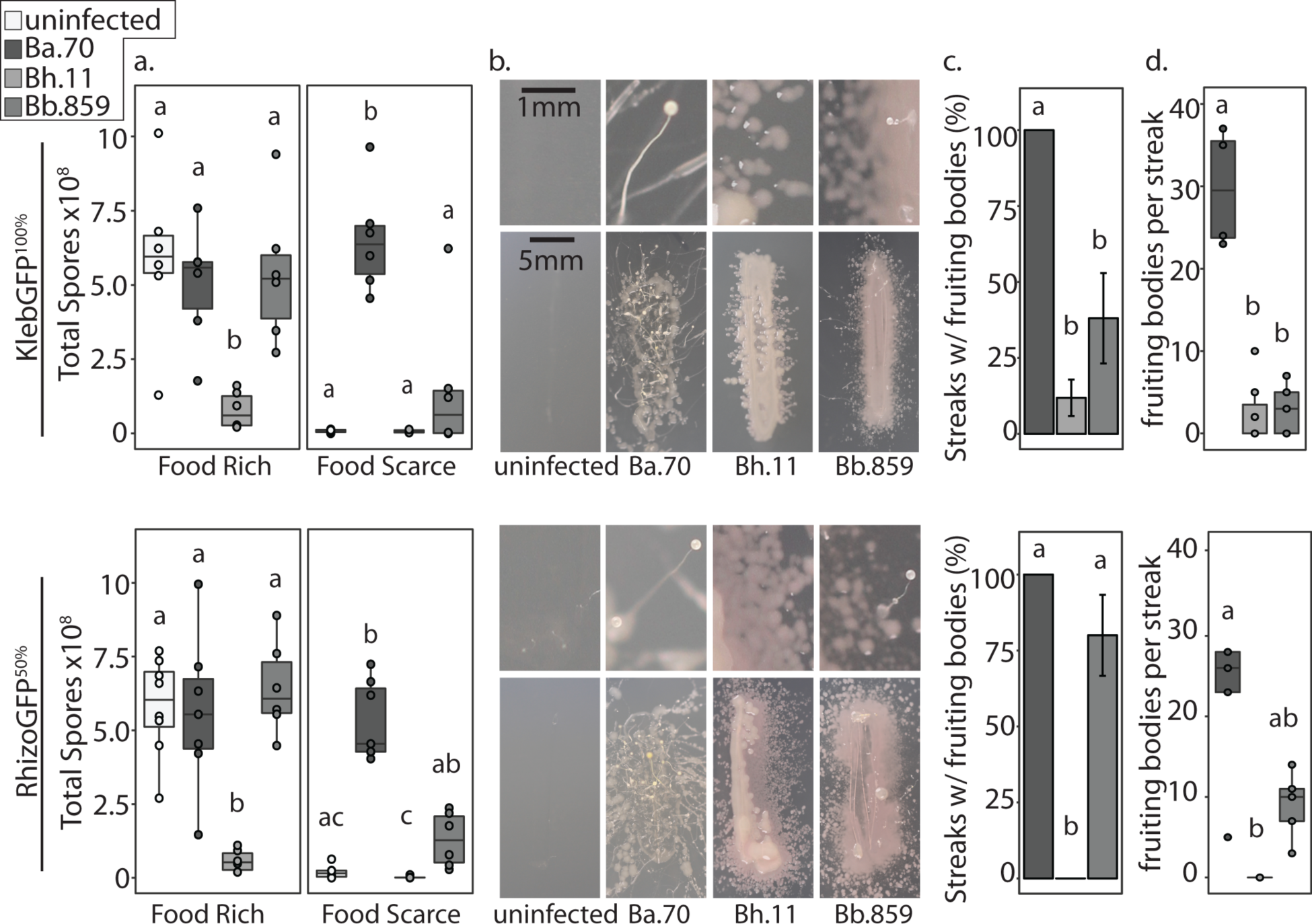
Farming Benefits Vary by Symbiont and Dispersal Strategy. For all figures, top panels represent data from sori pre-grown on *K. pneumoniae*-GFP^100%^ and bottom panels represent sori pre-grown on *Rhizobium*-GFP^50%^, both with (greys) and without (white) supplementation of 5% *Burkholderia.* a) Quantification of total spores harvested from food rich and food scarce plating conditions after transfer from *K.pneumoniae*-GFP^100%^ (top) or *Rhizobium*-GFP^5%^ (bottom) plating conditions. Points represent data from each individual replicate. b) Representative images of individual sori streaks from fruiting bodies developed on *K. pneumoniae*-GFP^l00%^ (top) or *Rhizobium*-GFP^50%^ (bottom) plates. Top panels are magnified sections of bottom panels. c) Percentage of bacterial positive sori streaks with observable fruiting bodies. Error bars ±SE. d) Number of fruiting bodies per fruiting body positive streaks for each individual replicate. All letters indicated post-hoc significance within panels. For streak tests, all sori where streaked five days after plating on *K. pneumoniae*-GFP^100%^ (top panels) or *Rhizobium*-GFP^50%^ (bottom panels) and all data was collected 5 days after streaking.

For the above assay, we evenly distributed spores on plates. If secondary bacteria are less numerous in host sori (as for *B. hayleyella* and *B. bonniea),* they might be spread too far from germinating spores to access and thus benefit from. Further, this assay might not best simulate spore dispersal in nature, where spores might be deposited in smaller denser patches by passing soil inhabitants. Therefore, we examined host fitness in food scarce conditions using a “streak” dispersal strategy. Here, we deposited individual sori from fruiting bodes grown on *K. pneumoniae*-GFP^100%^ or *Rhizobium*-GFP^50%^ in small patches (∼1-inch streaks) on nutrient medium. After a week of incubation, we measured the percent of fruiting body positive streaks and the number of fruiting bodies per each streak (Figure 5b-d).

Streaks from uninfected control sori did not produce bacterial colonies nor fruiting bodies. Over 95% of *Burkholderia* infected sori produced streaks with bacteria, however, the percent of streaks with fruiting bodies growing from these bacterial colonies significantly varies across *Burkholderia* species (x^2^=13.728 and 12.127, df=2, p=0.001 and 0.0023 for *K. pneumoniae*-GFP^100%^ and *Rhizobium*-GFP^50%^ conditions respectively). *Burkholderia* species also significantly influences the number of fruiting bodies per streak (x^2^=14 and 11.24, df=2, p<0.001 and 0.0036 for *K. pneumoniae-*GFP^100%^ and *Rhizobium*-GFP^50%^ conditions respectively). *B. agricolaris* infected sori generated significantly more fruiting bodies than *B. hayleyella* hosts from both conditions and *B. bonniea* hosts from *K. pneumoniae*-GFP^100%^ conditions. The number of fruiting bodies also increased the longer streak plates were left to incubate (Figure 6). Since fruiting bodies often developed from *B. bonniea* host sori, this suggests they gain better access to food under this dispersal strategy.

**Figure 6.**
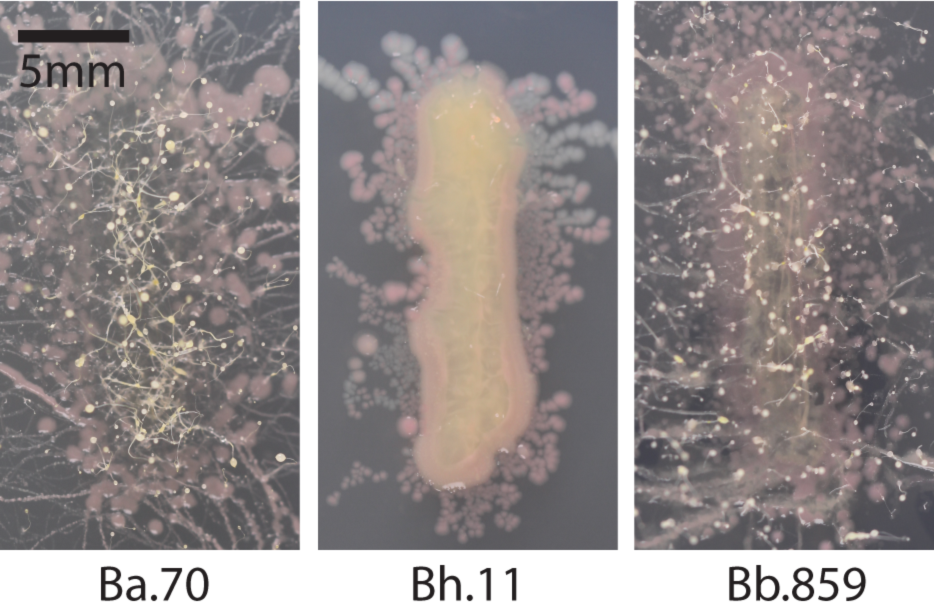
Fruiting Bodies Amplify in *Burkholderia* Infected Sori Streaks. Images of sori streaks two weeks after streak-testing. Sori were harvested from individual fruiting bodies that developed five days after plating spores on *Rhizobium*-GFP^50%^ culture conditions with 5% of the indicated *Burkholderia*-RFP strain.

We rarely witnessed fruiting bodies from *B. hayleyella* host sori streaks, providing an interesting counterpart to *B. bonniea.* Both symbionts induce similar densities of co-infection (Figure 4) yet differ in downstream benefits (Figure 5). This may be explained by the relative detriment each species exerts on its host. *B. hayleyella* reduces host fitness compared to *B. bonniea,* whereas *B. bonniea* hosts are often indistinguishable from uninfected counterparts (Figure 1 and 5). Thus, *B. hayleyella* toxicity may inhibit host development despite food availability.

### Co-infections and Conditional Benefits are Consistent Across *Burkholderia* Species Members

Our representative *Burkholderia* symbiont species significantly differed in their induction of co-infection and host impacts. We next asked whether these phenotypes were similar across strains of the same *Burkholderia* species. We imaged (Figure 7) and streak tested (Figure 8) host sori for additional *Burkholderia-RFP* strains after growth on *Rhizobium-GFP^50%^.* We again found noticeable, but low, levels of intracellular *Rhizobium* in *B. agricolaris* infected spores, with co-infection rare or absent in *B. hayleyella* and *B. bonniea* infected spores (Figure 5). The percent of spores infected by *Burkholderia* was significantly different depending on genotype (x^2^=22.65, df=5, and P<0.001), with *B. agricolaris* strains infecting fewer spores than *B. hayleyella* and *B. bonniea* (Figure 5b). Despite low levels of intracellular *Rhizobium* co-infection, we again frequently observed extracellular GFP in sori (Figure 7c).

**Figure 7.**
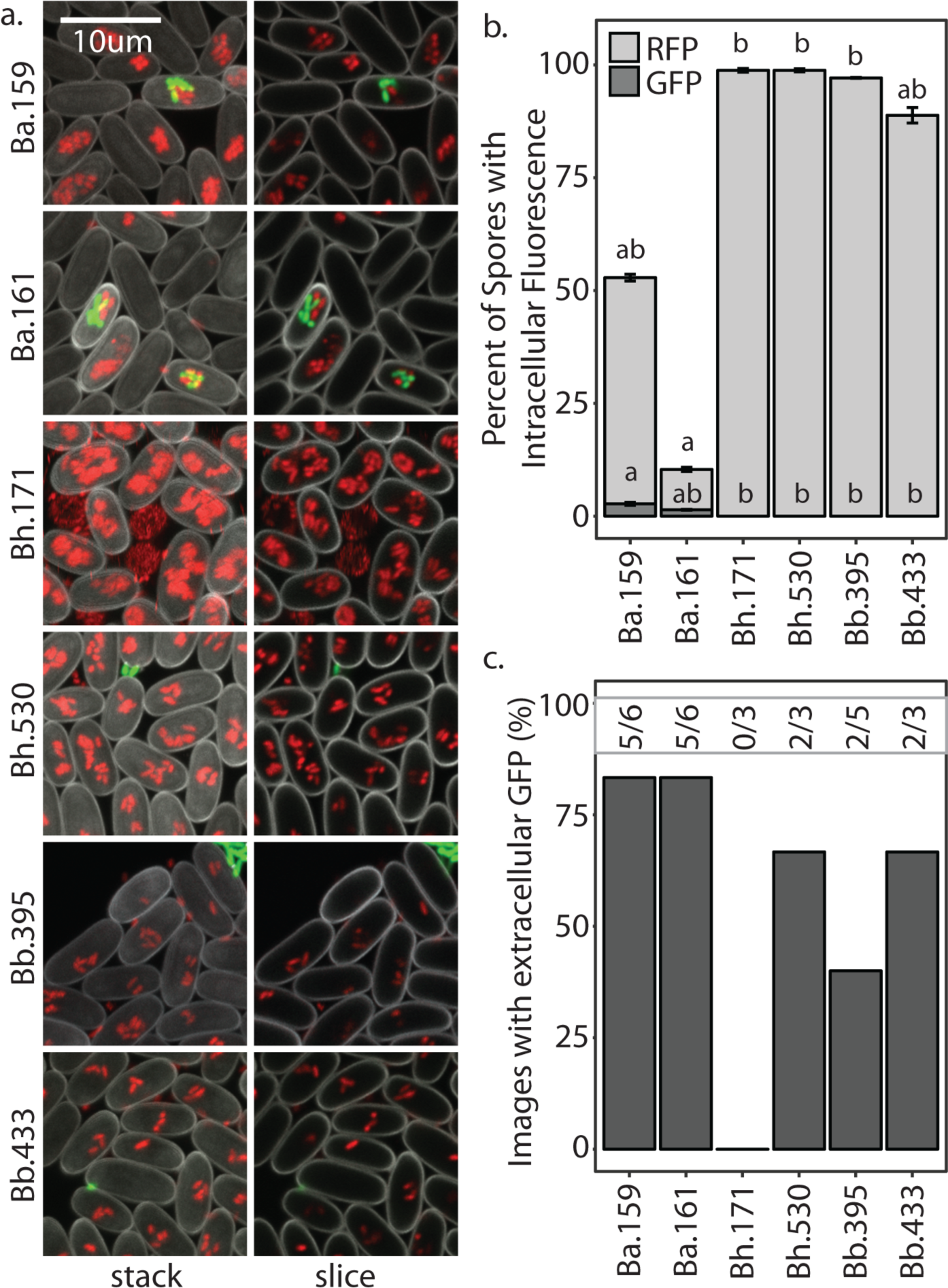
Co-infection Patterns Correspond to *Burkholderia* Species. **a)** Representative confocal micrographs of sori after five days post plating with 5% of the indicated *Burkholderia*-RFP strain and *Rhizobium*-GFP^50%^. b) Average percent of spores visualized with intracellular *Burkholderia*-RFP (light grey) and intracellular *Rhizobium*-GFP (dark grey) for each of the indicated *Burkholderia* species. Letters indicate post-hoc significance. c) Percent of images in which external GFP was visualized out of the total number of independent images acquired. Numbers above bars represent the raw number of external GFP positive images over the number of images acquired for each *Burkholderia* infection condition.

**Figure 8.**
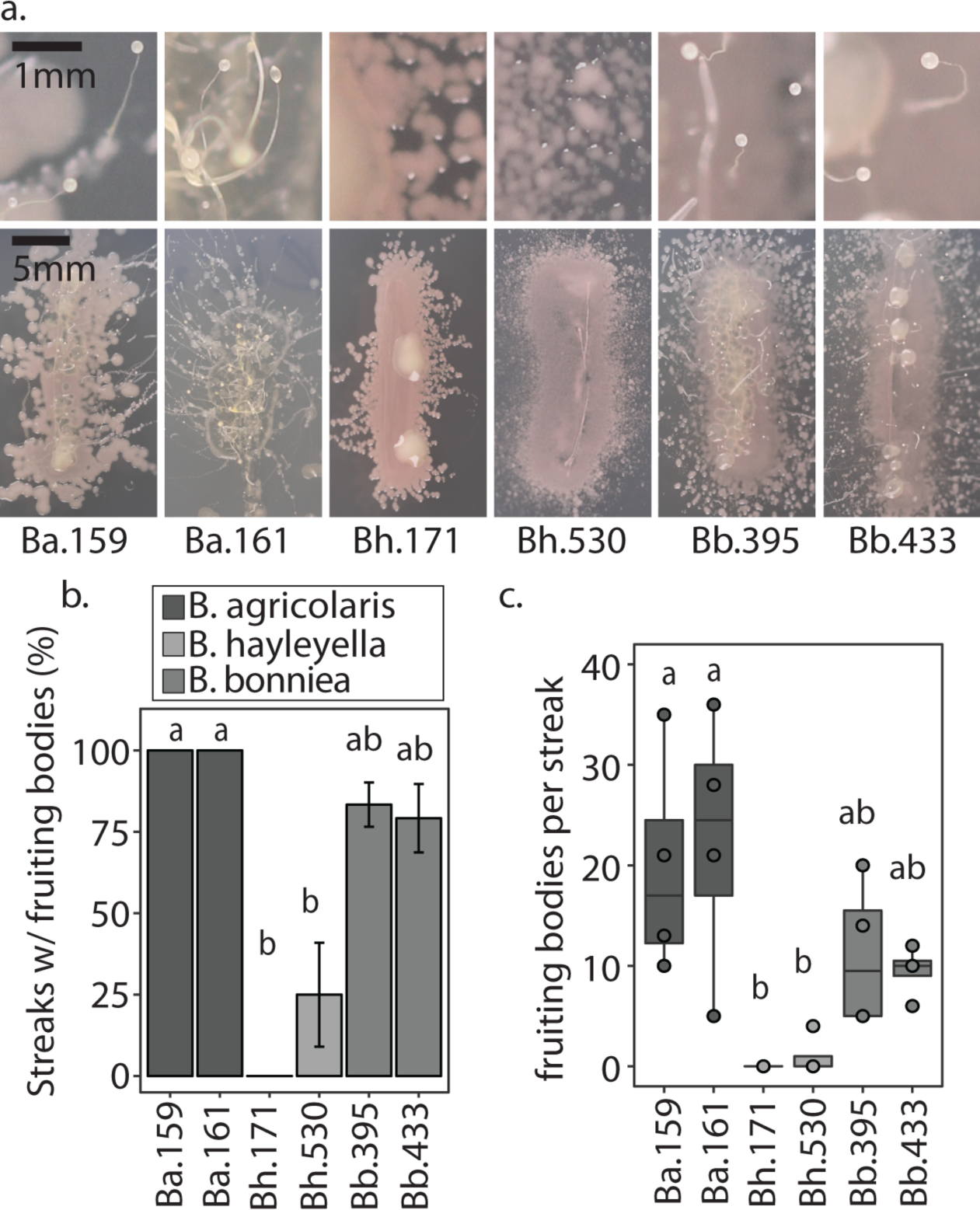
Host Benefits Correspond to *Burkholderia* Species and their Induction of Co-infections. **a)** Images of representative sori streaks from fruiting bodies grown on *Rhizobium*-GFP^50%^ with 5% of the indicated *Burkholderia* strain. Top panels are magnified sections of bottom panels. b) Percentage of bacterial positive sori streaks with observable fruiting bodies. Error bars ±SE. c) Number of fruiting bodies per fruiting body positive streaks for each individual. All letters indicated post-hoc significance within panels. All sori where streaked five days after plating on *Rhizibium*-GFP^50%^ and all data was collected 5 days after streaking.

Next, we investigated the benefits of infection by these strains in food scarce environments (Figure 8). We found that *Burkholderia* genotype significantly influences both the percentage of sori that generate fruiting bodies and the number of fruiting bodies in positive streaks (x^2^=12.127, df=2, p=0.002 and x^2^=1124, df=2, p=0.003, respectively). In accordance with our previous pattern *B. agricolaris* infections result in high, *B. bonniea* intermediate, and *B. hayleyella* low, levels of fruiting body production (Figure 8). These results demonstrate that species members similarly induce co-infections and result in similar subsequent host benefits.

## Discussion

Since elucidating the link between farming and *Burkholderia,* it has been assumed that all *Burkholderia* symbionts allow for secondary co-infections through an intracellular co-infection event. Intracellular co-infections of *B. agricolaris* with *K. pneumoniae* had previously been identified (34). Co-phagocytosis with *Burkholderia* and subsequent inhibition of phagocytic digestion was posited as a parsimonious mechanistic explanation for the phenomenon. *Burkholderia* can be visualized within intracellular vacuoles that appear similar to phagosomes (37) but how *Burkholderia* invades and survive within amoebae is not currently resolved. Nonetheless, in this scenario secondary bacteria must first be liberated from infected amoebae or spores (via regurgitation or host cell lysis) so that surrounding amoebae may reap the benefits of farming. Here, we found that co-intracellular infection is actually quite rare, occurring with some frequency for only *B. agricolaris* strains, and occurring differentially across secondary bacterial species. In contrast, all *Burkholderia* symbiont species generate extracellular secondary bacterial infection that can be visualized for all secondary species. The induction of overall secondary bacterial infection and its corresponding downstream benefits is significantly different across symbionts. *B. agricolaris* strains induce the highest levels of secondary infection (both intracellularly and overall) with *B. hayleyella* and *B. bonniea* strains generating exclusively extracellular infections at similarly low densities. This suggests that *Burkholderia* symbionts could induce secondary infections by multiple mechanisms that qualitatively and quantitatively differ between genotypes.

Overall, these results indicate that the predominant route by which different *Burkholderia* symbionts induce farming leans more towards extracellular bacterial carriage than intracellular co-infections. Susceptibility to secondary infections could be due to *Burkholderia* symbionts compromising the primitive immune system of their multicellular hosts. Sentinel cells serve as immune-like cells in multicellular slugs by trapping unwanted cargo through direct phagocytosis and/or neutralization by DNA net formation (31,32). When sentinel cells have accumulated cargo, they drop out the slug, thereby cleansing it of potential toxic entities (31). A gene deletion that reduces sentinel cells leads to retention of secondary bacteria through the slug stage and into the sorus (31). *Burkholderia* host slugs have fewer sentinel cells than uninfected counterparts and this defect goes away when hosts are cured of their *Burkholderia* symbiont via antibiotic treatment (44). Thus, the induction of secondary infections could be an indirect consequence of *Burkholderia* symbiosis resulting in sentinel cell reduction. This scenario may be comparable to the phenomenon of secondary infections in mammalian systems whereby primary infectious agents compromise the immune system of their hosts. Despite this, we cannot rule out the possibility that extracellular secondary infection originates from intracellular co-infection. It is possible that co-infected cells are more susceptible to lysis, rupturing and spewing their secondary bacterial passengers into the extracellular matrix. In either situation, secondary bacteria might then amplify within sori.

Intracellular co-infections are most frequent with *B. agricolaris* and the soil *Rhizobium* strain. This might reflect an ecologically relevant association between these species in nature. *Burkholderia* and *Rhizobium* are both ubiquitous in soil and contain several important symbiont species which have been found co-colonizing the same hosts (45–48). Predation by amoebae in soil and aquatic systems shapes microbial community assembly and overall food webs (49). Given the likely co-occurrence of soil amoebae with *Burkholderia* and other soil microbes it’s tempting to speculate on how these multipartite interactions influence overall microbial communities and higher trophic levels. Here, we show that amoebae co-disperse *Burkholderia* symbionts and secondary bacterial hitchhikers to new environments. Thus, the impact of amoebae on their surrounding microbial network can go well beyond predator-prey dynamics. Finally, our observation of *Burkholderia* and *P. aeruginosa* co-infection amplifies the concern that soil amoebae can serve as reservoirs for bacterial pathogens. These results suggest that *Burkholderia* symbionts can increase the suite of potential pathogenic partners hosted by amoebae.

*Burkholderia-fungal* associations have been well recognized for their importance in the soil ecosystem and for their bio-restoration potential (50–52). There are compelling parallels between *Burkholderia-Dictyostelium* and *Burkholderia-fungal* associations. Some *Burkholderia* (notably *B. terrae*) are capable of adhering to and migrating with growing fungal hyphae through soil (53). Similar to our system, some of these fungal associates assist in the co-migration of other (non-migrating) bacteria (54). Several mechanisms have been proposed to underlie these interactions, such as direct receptor binding and indirect biofilm co-aggregation (55). *B. terrae* extracellularly colonizes fungal hyphae but many other *Burkholderia* symbionts of diverse hots persist intracellularly (13,56). A particularly interesting example is the *Rhizopus microsporus* endosymbiont *B. rhizoxinica* fungi, which produces the rice seedling blight toxin (57). Recently, secretion systems have been shown to be important for the active invasion of *B. rhizoxinica* across the fungal cell wall and into the host cytoplasm (56). Secretion systems have also been implicated in *B. psuedomallei* infections (58,59). However many plant mutualistic *Burkholderia* species, which are closer relatives to *Burkholderia* symbionts of *Dictyostelium,* appear to lack some of these systems (58–60)(58–60). The hypothesized portal of entry into *Dictyostelium* is via phagocytosis, which could circumvent the need for invasion specific mechanisms. Overall, *Burkholderia* symbionts of other hosts can help inform our understanding of the *Burkholderia-amoebae* symbiosis and vice versa.

Biofilm formation is particularly intriguing to consider as a mechanistic explanation of secondary infection. *Burkholderia* adherence to secondary bacteria would increase the likelihood of co-phagocytosis or extracellular co-colonization. Different adhesive capacities of *Burkholderia* and secondary species could explain differences in the extent of secondary infections across bacterial combinations. Indeed, co-aggregation could be a simple explanation for the observation of high levels of co-infection with *B. agricolaris* and *Rhizobium.* Interestingly, recent work implicates *Dictyostelium* lectins in the farming phenomenon, higher lectin expression was detected in farmer *D. discoideum* clones and addition of endogenous lectins induced bacterial carriage (61). Although this work did not consider the presence or impact of *Burkholderia,* we think *Burkholderia* symbionts play a key role. *Burkholderia* could induce farming via induction of lectin expression in amoebae or more simplistically, *Burkholderia* lectins may mediate adherence of secondary bacteria. Indeed, lectin expression by *B. cenocepacia* is an important component of biofilm formation and lectin aids in adherence of *B. cepacia* to host tissues (42,62). Future exploration into lectin expression and adhesion mechanisms will be helpful for clarifying these themes.

In addition to elucidating the phenomenon of secondary infections our results exemplify the context dependency of symbiotic outcomes in this system. We found that the costs and benefits of this symbiosis can be modified by different bacterial conditions and spore dispersal processes. The nature and extent of farming induction by *Burkholderia* symbionts differs across symbiont species and so do their corresponding contextual fitness outcomes. Ultimately, further research into the mechanisms, consequences, and ecological framework of the *Burkholderia-Dictyostelium* symbiosis will help illuminate microbial interaction dynamics relevant to infection biology and microbial ecology.

## Acknowledgements

We thank Kyle Skottke for fruitful discussions and manuscript review, Joan Strassmann and David Queller for initial guidance in the system, and all members of the DiSalvo lab at SIUE, particularly Jacob Miller for help with experimental discussions.

## Conflict of interest

We declare no conflict of interests.

## Author Contributions Statement

NK and SD designed the study. NK and SD performed experiments with assistance from ME, RN, and TH. SD and TH wrote the manuscript.

## Funding

This study was supported by SIUE start-up funds from the DiSalvo lab.

## Data Availability Statement

The raw data supporting the conclusions of this manuscript will be made available by the authors, without undue reservation, upon request.

